# Flexible brain state engagement predicts cognitive control transdiagnostically

**DOI:** 10.64898/2025.11.30.691395

**Authors:** Jean Ye, Saloni Mehta, Milana Khaitova, Jagriti Arora, Fuyuze Tokoglu, C. Alice Hahn, Cheryl Lacadie, Abigail S. Greene, Marisa N. Spann, Gerard Sanacora, Scott W. Woods, Vinod H. Srihari, R. Todd Constable, Dustin Scheinost

## Abstract

Cognitive control supports adaptive responses in an ever-changing world. While alterations in cognitive control have been consistently observed in a range of psychiatric disorders, the neural mechanisms giving rise to this behavioral variation remain elusive. Here, we tested whether the ability to flexibly recruit recurring brain activation patterns (i.e., brain states) may serve as an intermediate phenotype supporting cognitive control in individuals with a spectrum of clinical symptoms. We leveraged machine learning and external validation to explore this question in three independent, transdiagnostic datasets (N>600), including participants with anxiety disorders, schizophrenia, mood disorders, substance use disorders, post-traumatic stress disorder, obsessive-compulsive disorder, and neurodevelopmental disorders. To capture cognitive control’s multifaceted nature, we assessed two of its components—inhibition and shift— using both task-based and questionnaire data. Flexible brain state engagement predicted all cognitive control metrics in previously unseen individuals transdiagnostically, regardless of which dataset was used for model training. Connectome-based predictive modeling also revealed that shared brain networks underpinned flexible brain state engagement in a transdiagnostic manner. Leveraging brain network dynamics, we further observed that moments of more flexible brain state engagement aligned with moments of network connectivity related to better cognitive control within the same individual. This temporal alignment was replicated in all three datasets with heterogeneous samples. Altogether, this study suggests flexible engagement of brain states may support both inter- and intra-individual differences in cognitive control across individuals with diverse mental health profiles.

**Funding sources:** This work was supported by F31AA032179 and R01MH121095. Data were provided in part by the Consortium for Neuropsychiatric Phenomics (NIH Roadmap for Medical Research grants UL1-DE019580, RL1MH083268, RL1MH083269, RL1DA024853, RL1MH083270, RL1LM009833, PL1MH083271, and PL1NS062410) and the Transdiagnostic Connectome Project (TCP) dataset. Datasets were accessed through OpenNeuro (https://openneuro.org/datasets/ds005237 and https://openneuro.org/datasets/ds000030/versions/00016).

## Introduction

Cognitive control plays a crucial role in our navigation through life^1^. It supports responses to the rich influx of information we receive every moment. For instance, during a conversation, cognitive control allows us to ignore an incoming text notification while directing our attention to a loud alarm sounding in the room. As demonstrated by this example, cognitive control is a multifaceted construct^2,3^, encompassing both the ability to focus on the current task in the face of distraction (i.e., inhibition) and the flexibility to switch tasks when needed (i.e., shift).

There are remarkable variations in cognitive control across individuals. Notably, alterations have been consistently observed across psychiatric conditions, including schizophrenia^4^, mood^5^, substance use^5^, obsessive-compulsive (OCD)^6–8^, and neurodevelopmental disorders^9–11^. Given cognitive control’s central role in our daily life, understanding how variations in brain responses contribute to individual differences in cognitive control is essential. Additionally, as aberrant cognitive control cuts across psychiatric disorders^12^, studying it transdiagnostically allows for the consideration of a broad spectrum of behavioral variations. This approach also aligns with recent efforts, such as the Research Domain Criteria (RDoC) framework^13,14^, to consider brain-behavior relationships on a continuum.

As cognitive control involves dynamic responses to a changing environment, such effective behaviors may require corresponding neural dynamics^15^. Tracking moment-to-moment fluctuations in brain activity is now feasible, as new advances identify brain states (e.g., recurring patterns of brain activation or connectivity) to examine their engagement over time using functional magnetic resonance imaging (fMRI)^16–18^. Furthermore, recent literature has noted that while once considered a source of noise, variations in brain signals (i.e., neural variability) may confer flexibility in exploring and recruiting brain network configurations in response to evolving demands^9,19,20^. Thus, both brain state dynamics and neural variability are appealing candidate mechanisms for cognitive control. While emerging work has begun to investigate how brain state dynamics or neural variability supports cognition^21–26^, there is a paucity of work studying whether these two brain metrics jointly predict individual differences in cognitive control. How these associations unfold transdiagnostically remains even more unclear.

Here, we address this gap by examining the relationship between cognitive control and a well-validated metric that combines brain state dynamics and neural variability. State engagement variability (SEV)^27^ quantifies variability in continuous brain state engagement to approximate how flexibly individuals recruit recurring brain states. Individuals with psychopathology have shown lower SEV than healthy controls (HCs)^27,28^, and lower SEV has been linked to worse executive function during development^29^. To test if SEV supports cognitive control transdiagnostically, we leveraged three independent datasets recruiting individuals with a range of clinical diagnoses, including schizophrenia, mood, anxiety, bipolar, substance use, and neurodevelopmental disorders. To consider cognitive control’s multifaceted nature, we included measures to capture two of its aspects, inhibition and shift. The association between cognitive control and SEV was established in two ways. First, external validation assessed whether SEV predicts cognitive control in previously unseen individuals. Secondly, we used network dynamics to investigate whether moment-to-moment fluctuations in SEV and cognitive control network dynamics aligned over time. As a secondary analysis, we also examined group differences in the relationship between SEV and cognitive control.

## Results

We probed the relationship between SEV and cognitive control in three transdiagnostic datasets. First, we treated the YaleNeuroConnect dataset^30^ as our main dataset (92 HCs; 145 patients). This richly phenotyped dataset recruited the largest number of patients with a range of psychiatric disorders and comorbidities, providing a great resource to study brain-behavior associations transdiagnostically. Here, inhibition was evaluated using a metric from the Delis-Kaplan Executive Function System battery, specifically the scaled scores from the color-word incongruent condition. We focused on the incongruent condition since metrics from one task condition have demonstrated greater reliability and validity than a difference score^31–36^. Shift was assessed using the shift subscale (normed T score) from the Behavioral Rating Inventory of Executive Function (BRIEF).

We then validated our findings externally in two independent datasets. To validate inhibition findings, we selected the UCLA Consortium for Neuropsychiatric Phenomics dataset^37^ (109 HCs; 116 patients; hence referred to as the inhibition validation dataset). This dataset recruited HCs and individuals with schizophrenia, bipolar disorder, or attention-deficit/hyperactive disorder (ADHD). Inhibition was measured using Stroop incongruent mean response time (RT; in milliseconds).

Shift results were externally validated in the Transdiagnostic Connectome Project dataset^38^ (69 HCs; 94 patients; hence referred to as the shift validation dataset), which, similar to the main dataset, also recruited individuals with various psychiatric diagnoses. To best match the main dataset’s shift item, we used the Rumination Responsiveness Scale (RRS) to assess flexibility in thinking style. All behavioral measures were collected outside the scanner. For all behavioral metrics, a higher score indicated greater difficulty with inhibition or shift.

### SEV correlated with cognitive control within datasets

We first established a link between SEV and cognitive control within datasets. SEV was computed following an established framework^27^ (**Supplementary Figure 1**). In brief, we first replicated prior work^39^ to identify four recurring brain states in the Human Connectome Project^40^ (HCP; **Supplementary Materials**) dataset. We next tracked their continuous engagement in the transdiagnostic cohorts. Brain states were generated in HCP for two reasons. Identifying brain states in an independent dataset circumvents circular analysis. Secondly, HCP included a range of tasks, allowing the extraction of brain states supporting low-(e.g., motor movement) to high-level (e.g., working memory and emotion processing) behaviors. Indeed, four distinct but recurring brain states associated with different task conditions were found using nonlinear manifold learning^39^. Based on the task conditions linked to each brain state, we labeled them as fixation, high-cognition, low-cognition, and transition (**Supplementary Table 1**). Previous work suggested these brain states also appeared during rest^39^. Additionally, our framework allows both spatial and temporal overlap in brain state engagement. Such flexibility permits the simultaneous combination of multiple brain states at each time point, aligning with the possibility that multiple cognitive processes may be active during rest.

While HCP mainly recruited healthy adults, prior literature suggests that similar brain states can be found in HCs and patients^39,41,42^. However, for further validation, we repeated our analyses after generating brain states from a transdiagnostic cohort and obtained consistent results (**Supplementary Materials**).

SEV was defined as the standard deviation of continuous brain state engagement and was extracted from all three transdiagnostic cohorts’ resting-state fMRI scans. As the purpose of this analysis was to establish a link between SEV and cognitive control before prediction, we limited the number of tests run by extracting an overall SEV measure with principal component analysis (PCA). The first component accounted for 92.91%, 93.34%, and 90.29% of the variance in the main, inhibition validation, and shift validation datasets, respectively. Overall SEV may thus reflect a general ability to flexibly recruit and combine recurring brain states as various demands arise.

For inhibition in the main dataset, we found that lower SEV was associated with worse inhibition in patient participants (r=-0.18, p=0.03; **Figure 1**) but not in HCs (r=0.05, p=0.63). In the inhibition validation dataset, lower SEV was associated with worse inhibition across all participants (r=-0.27, p < 0.01; **Figure 1**). However, a post-hoc analysis in the inhibition validation dataset revealed a significant group interaction (z=-2.43, p=0.02; **Supplementary Table 3**). Altogether, findings from both the main and inhibition validation datasets suggested that patients drove the association between SEV and inhibition.

**Figure 1.**
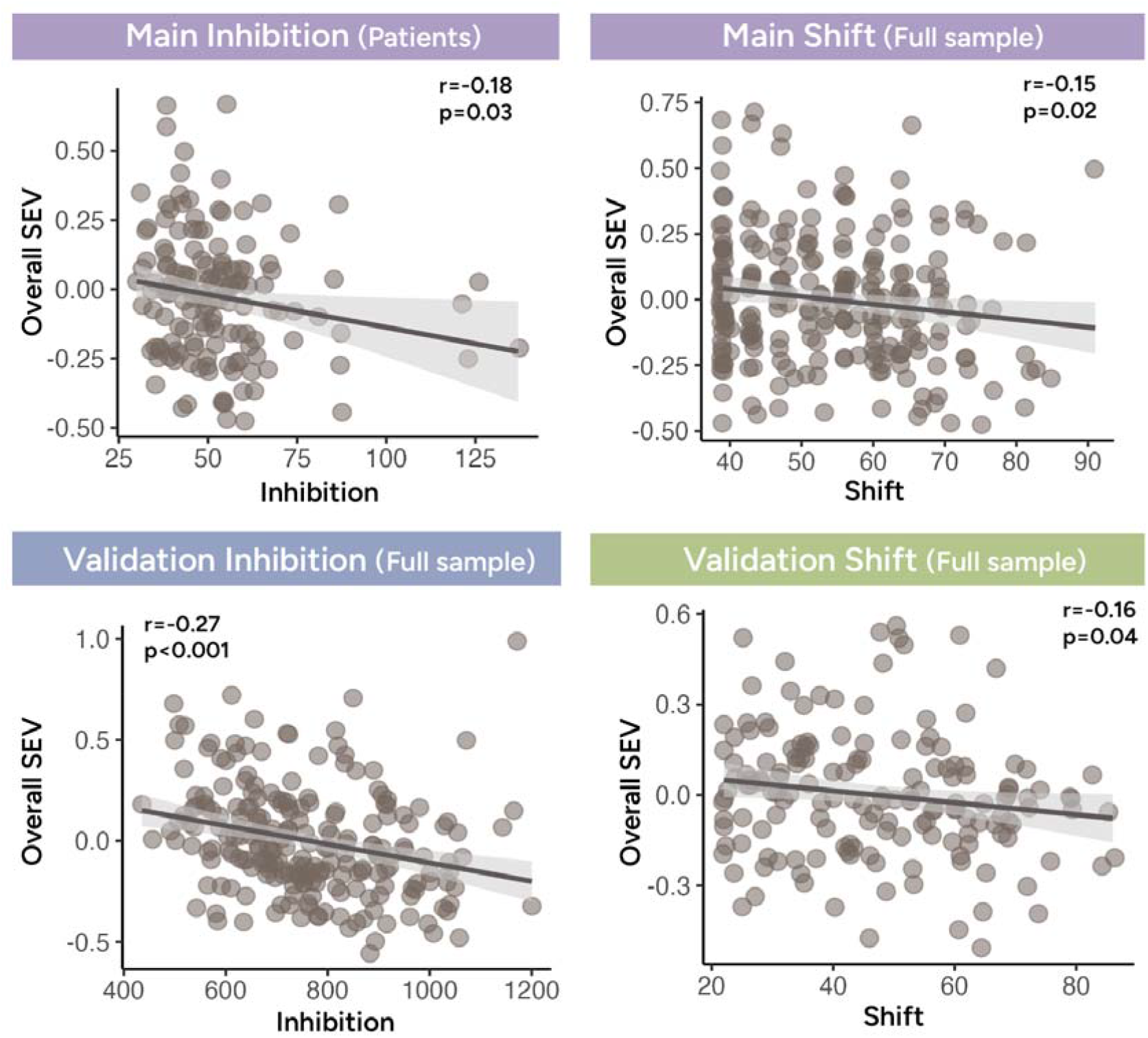
Within-dataset relationships between SEV and cognitive control. We found that lower SEV, indicating reduced flexibility in brain state engagement, was correlated with worse out-of-scanner task-based and questionnaire measures of cognitive control across all three transdiagnostic datasets.

Lower SEV was additionally associated with worse shift in all participants from the main dataset (r=-0.15, p=0.02; **Figure 1; Supplementary Table 4**) and all participants from the shift validation dataset (r=-0.16, p=0.04; **Figure 1; Supplementary Table 4**). Across various clinical conditions, lower SEV correlated with worse cognitive control. This pattern aligned with the interpretation that lower variability may reflect reduced flexibility in recruiting brain states to address evolving demands. Importantly, these findings lay the foundation for using machine learning to predict cognitive control from SEV in external samples.

### SEV predicted cognitive control in external samples

Next, we investigated whether we could train a linear model to predict cognitive control from SEV in previously unseen individuals. We first trained a model in the main dataset before testing it in the validation datasets. To further demonstrate the robustness of our results, we reversed the datasets before repeating the analysis (i.e., trained on the validation datasets before testing on the main dataset). Model performance was evaluated by correlating predicted and observed cognitive control.

Models were built using SEVs from all four brain states. A linear model was first trained to predict inhibition in the main dataset before being applied to the inhibition validation dataset (**Figure 2A**; model parameters in **Table S5**). Since we previously noted that patients may drive the relationship between SEV and inhibition, model training and testing were conducted only with patient participants. This model successfully predicted inhibition in the inhibition validation dataset (r=0.32, p<0.001; **Figure 2B**). Next, we switched the dataset and found that a newly trained model in the inhibition validation dataset also successfully predicted inhibition in the main dataset (r=0.20, p=0.02; **Figure 2B**; model parameters in **Table S5**).

**Figure 2.**
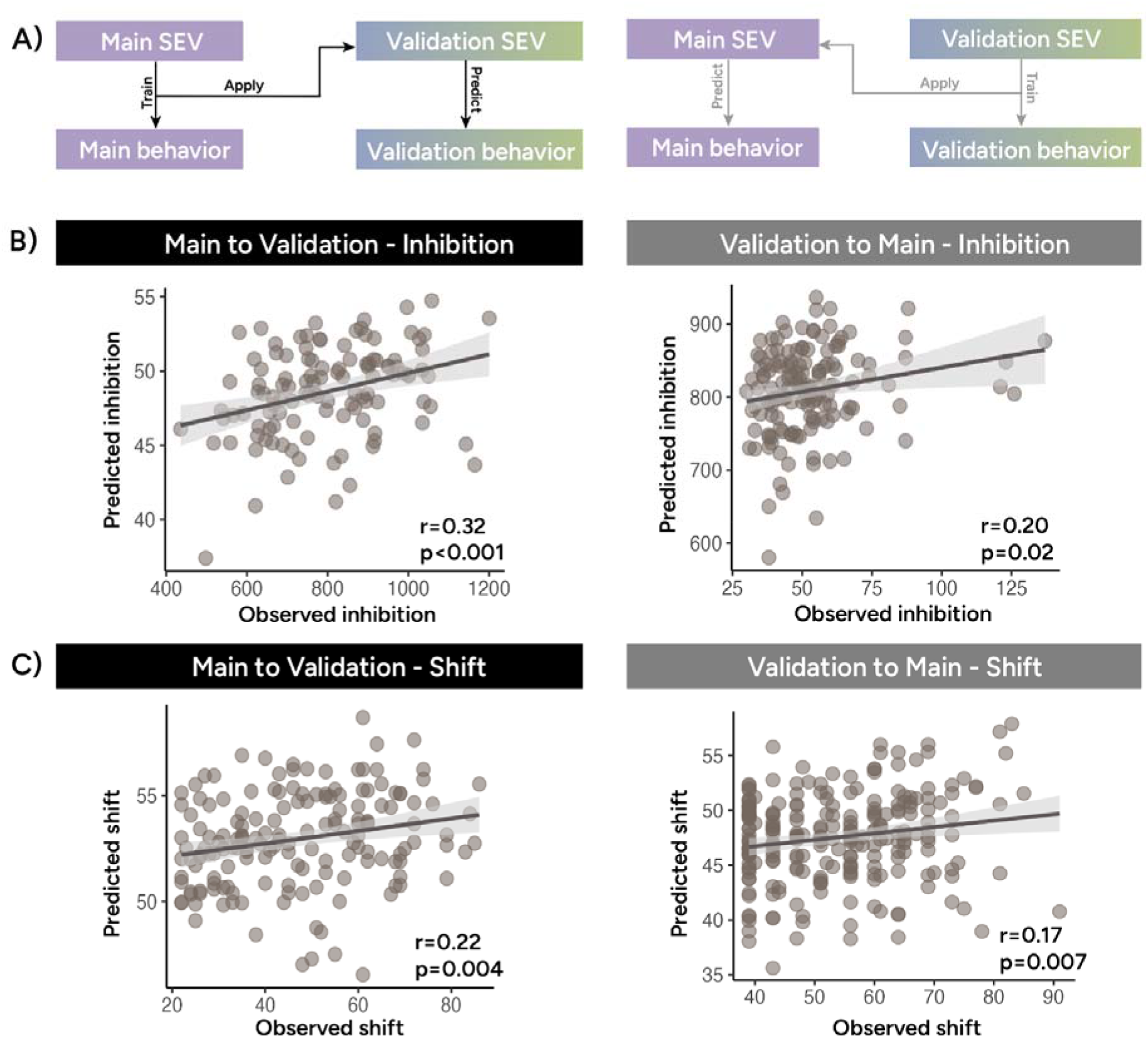
SEV predicted cognitive control in external samples. **A)** Linear models were trained to predict cognitive control using SEV in the main dataset before the models were applied to the validation datasets. We also performed the analysis again after swapping the datasets (i.e., training on one of the validation datasets before testing on the main dataset). We found that SEV can successfully predict both inhibition **B)** and shift **C)** in previously unseen individuals, regardless of which sample was used for model training.

For shift, since the relationship between SEV and shift appeared in the full sample, all participants from the main and shift validation datasets were used for training and testing. A model trained in the main dataset successfully predicted shift in the shift validation dataset (r=0.22, p=0.004; **Figure 2C**; model parameters in **Table S5**). After switching the datasets, we again found that a new model trained in the shift validation dataset can also predict shift in the main dataset (r=0.17, p=0.007; **Figure 2C**; model parameters in **Table S5**). Successful external validation demonstrated that regardless of the dataset used for model training, SEV can predict various metrics of cognitive control in previously unseen individuals. Importantly, consistent results were obtained using SEV generated from brain states identified in a transdiagnostic cohort (**Supplementary Materials**).

### Shared brain networks supported SEV

Given its crucial role in supporting cognitive control, we next asked which brain networks support SEV. To this end, we leveraged 10-fold connectome-based predictive modeling^43^ (CPM) to predict resting-state SEV using functional connectivity (FC) from an in-scanner cognitive control paradigm in each dataset (main and inhibition validation datasets: stop signal task; shift validation dataset: Stroop task). We opted to use FC from a cognitive control task for two main reasons. First, since the previous section demonstrated a close link between SEV and cognitive control, FC during a cognitive control task may be relevant for revealing the brain networks supporting SEV. Secondly, we alleviated concerns about data leakage and circular analyses by using FC from task-based fMRI to predict resting-state SEV. As above, model performance was assessed by correlating predicted and observed SEV, with significance determined by permutation testing. Only edges that significantly predicted SEV across all 10 folds were included in the final SEV network. This step also returned a positive (where greater connections predicted higher SEV; shown in red in **Figure 3A-C**) and negative SEV networks (where greater connections predicted lower SEV; shown in blue in **Figure 3A-C**).

**Figure 3.**
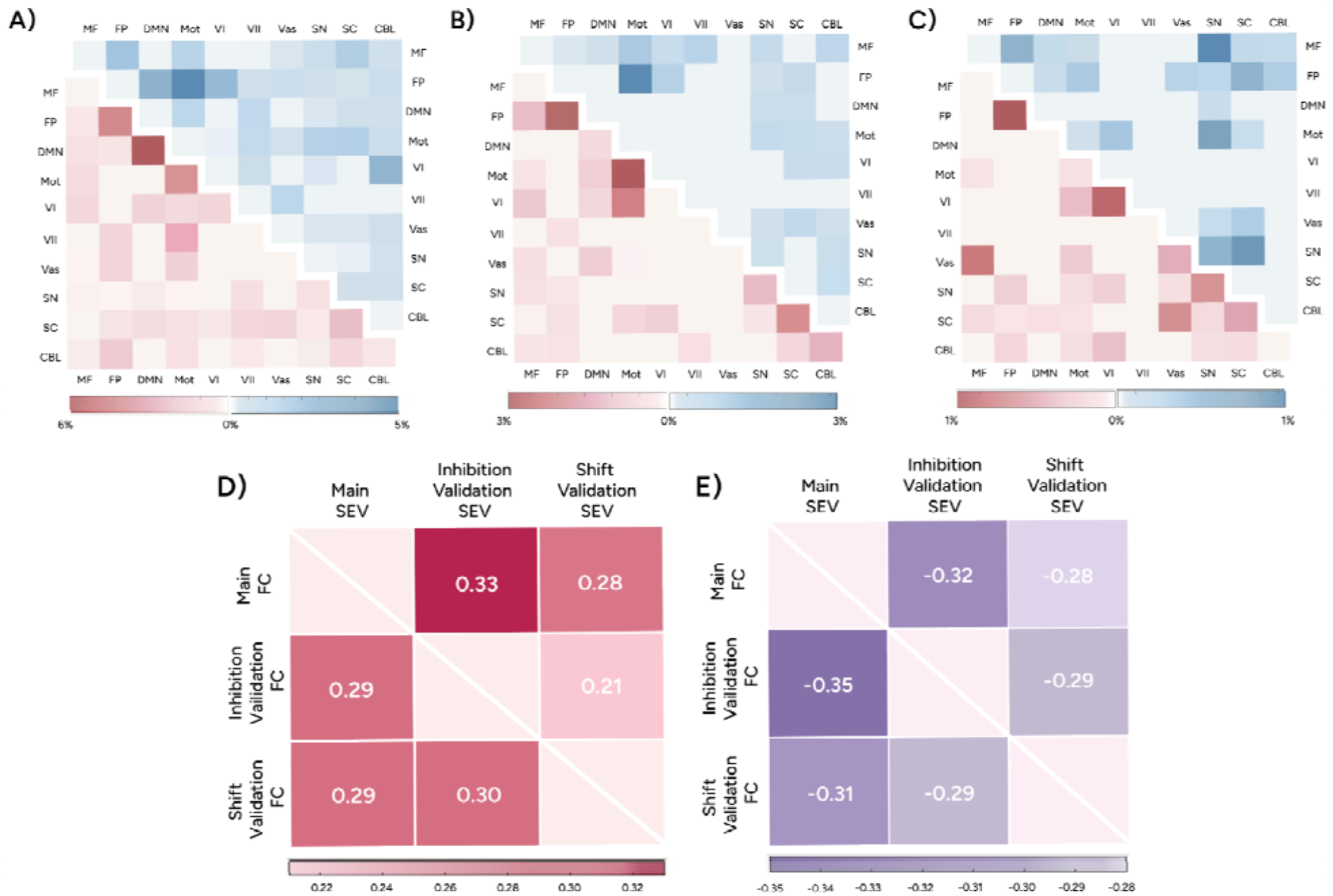
Brain networks underpinning flexible engagement of brain states. We identified brain networks supporting SEV in main **A)**, inhibition validation **B)**, and shift validation **C)** datasets using 10-fold connectome-based predictive modeling. The lower triangular portion (red) indicated the positive SEV network, whereas the upper triangular part (blue) indicated the negative SEV network. We normalized the number of significant edges by the total network size (i.e., the number of edges between or within networks). Both the SEV positive **D)** and negative **E)** networks predicted resting-state SEV in the other two datasets. Correlation values between SEV network functional connectivity and SEV are shown in matrices. MF, medial frontal network; FP, frontoparietal network; DMN, default mode network; Mot, motor network; VI, visual I network; VII, visual II network; VAs: visual association network; SN, salience network; SC, subcortical network; CBL, cerebellum; FC: functional connectivity; SEV: state engagement variability.

We were able to predict overall SEV in all three datasets transdiagnostically (main: r=0.37, p=0.002; inhibition validation: r=0.32, p=0.004; shift validation: r=0.28, p=0.002). A closer look at these SEV networks revealed that across datasets, within-network connections predicted higher SEV (**Figure 3A-C**). In contrast, lower SEV was predicted by stronger FC between the motor and various cognitive control networks (**Figure 3A-C**).

While all three SEV networks exhibited spatial overlap (ps<0.05; **Supplementary Tables 6 & 7**), some variations existed (**Figure 3A-C**). This is likely because different cognitive control tasks (i.e., stop signal vs. Stroop) were used to identify SEV networks across datasets, contributing to different edges being included in each network. In support of this speculation, the SEV network identified in the shift validation dataset (which collected Stroop) appeared more visually distinct from the SEV networks identified in the main and inhibition validation datasets (both collected stop signal; **Figure 3A-C**). Thus, we tested whether each SEV network generalized across the transdiagnostic datasets. Specifically, we computed each SEV network’s FC during task-based fMRI in the two external datasets. SEV network FC was then correlated with resting-state SEV in each external dataset to interrogate network generalizability. Notably, all three SEV networks successfully predicted SEV in any external dataset (**Figures 3D-E**). Altogether, these results strongly indicated that shared brain networks support flexible engagement of brain states across multiple datasets in a transdiagnostic manner.

### Moment-to-moment changes in SEV and cognitive control networks aligned over time

The previous section established that SEV can predict inter-individual differences in cognitive control. Is SEV also sensitive to fluctuations in cognitive control within the same person? Studying their continuous interaction was challenging when we did not have access to instantaneous cognitive control or SEV across the three datasets. Cognitive control is often evaluated using information drawn from seconds-long task trials. On the other hand, SEV computation requires tracking brain state engagement over an extended period of time. These challenges make it difficult to study how cognitive control and SEV fluctuate from one moment to another in a continuous manner. Here, we bypassed these roadblocks by adopting a novel approach. Specifically, we computed instantaneous network engagement to predict moment-to-moment changes in SEV and cognitive control to investigate their alignment over time.

As this approach required brain networks predictive of each behavior of interest, we used the main dataset, which recruited the largest number of participants, to identify brain networks predicting cognitive control, transdiagnostically. Using 10-fold CPM and FC from a stop signal paradigm, we successfully identified an inhibition (r=0.27, p<0.001; **Figure 4A**) and a shift network (r=0.25, p<0.001; **Figure 4A**). Similar to above, we only retained edges that predicted cognitive control across all 10 folds in the final cognitive control networks.

**Figure 4.**
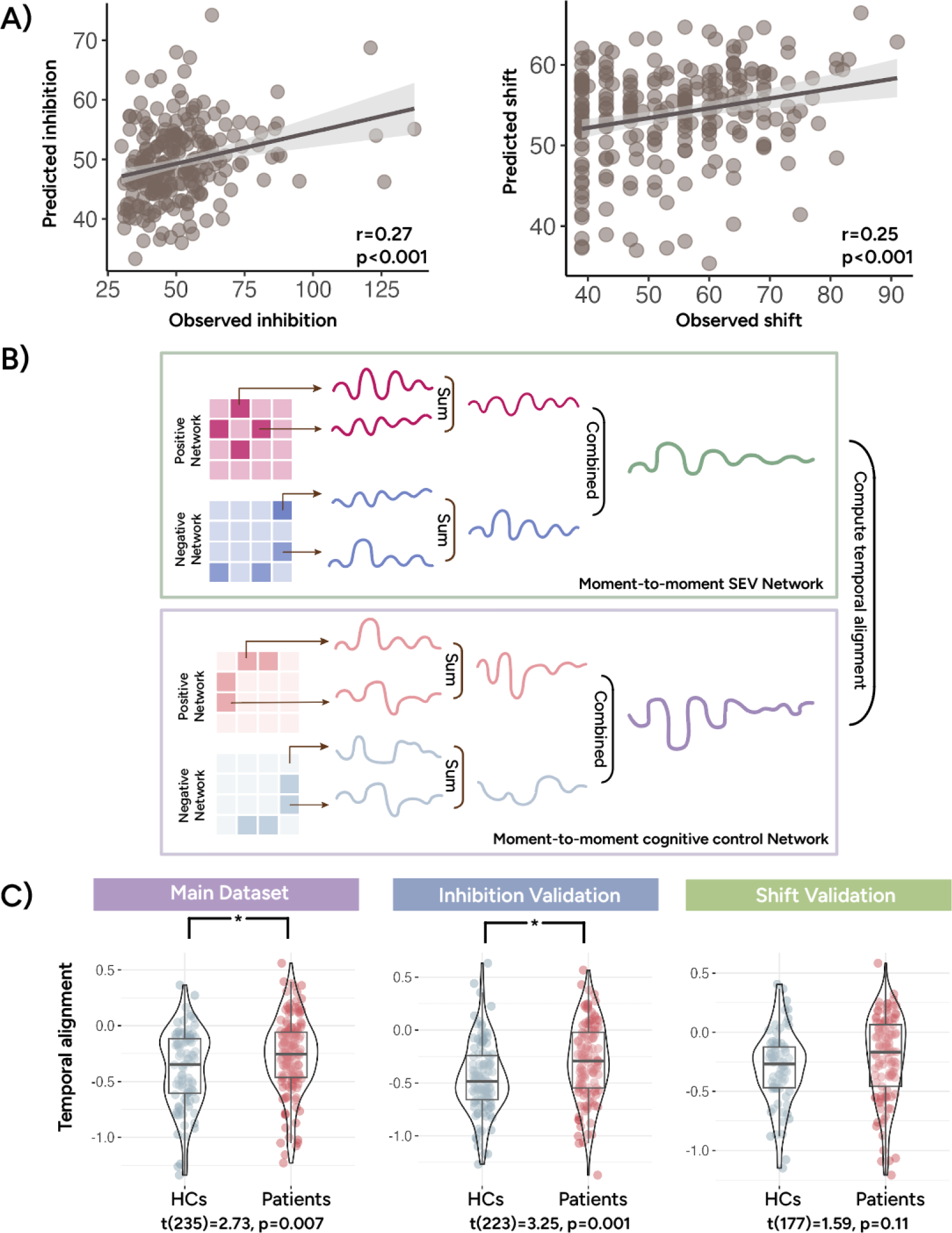
Moment-to-moment changes in SEV and cognitive control networks aligned over time. In our main dataset, we successfully identified brain networks predicting inhibition and shift **A)**. Using these brain networks, we investigated if moment-to-moment engagement of the SEV network aligned with moment-to-moment engagement of the cognitive control network **B)**. Moments of higher SEV corresponded to moments of functional connectivity related to better cognitive control. We further found that the temporal alignment between SEV and shift network dynamics was stronger in HCs than in patients in the main and inhibition validation datasets **C)**.

We next applied edge time series to the SEV network (**Figure 3**) and cognitive control network (**Figure 4A**) to track their instantaneous engagement during resting-state fMRI. Edge time series^44,45^ is a recently introduced technique that can decompose FC between two brain regions into its moment-to-moment contribution. It is computed by finding the element-wise product of the activation time series from a pair of brain regions after z-scoring. Higher values in this co-activation timeseries reflect moments where the two brain regions activate in the same direction. Following a prior framework^46^ (see illustration in **Figure 4B**), we computed edge time series for each edge in each positive or negative CPM network before summing across edges to assess overall moment-to-moment positive or negative network engagement. Finally, negative network engagement was subtracted from positive network engagement to evaluate the continuous engagement of the whole CPM network.

Tracking each CPM network’s continuous engagement allowed us to predict and approximate how SEV and cognitive control fluctuated over time. Importantly, leveraging network dynamics to estimate SEV and cognitive control provided the opportunity to track their alignment instantaneously. By correlating these networks’ moment-to-moment engagement, we estimated whether the two constructs fluctuated continuously within the same individual (**Figure 4B**). One-sample t-tests were performed to examine temporal alignment (i.e., if correlation values differed significantly from 0). Across all three datasets, we found a strong temporal alignment between fluctuations in SEV and cognitive control network dynamics. Specifically, brain network dynamics revealed that moments of greater SEV network connectivity occurred alongside moments of network connectivity related to better cognitive control (SEV and inhibition networks: t(236)=-22.92, p<0.001; SEV and shift networks: t(236)=-14.07, p<0.001). Notably, we externally replicated this pattern in both the inhibition validation dataset (SEV and inhibition networks: t(224)=-15.39, p<0.001; SEV and shift networks: t(224)=-15.32, p<0.001) and shift validation dataset (SEV and inhibition networks: t(178)=-20.27, p<0.001; SEV and shift networks: t(178)=-10.04, p<0.001).

In a post-hoc analysis, we further probed group differences in the temporal alignment between SEV and cognitive control network dynamics. Alignment between inhibition and SEV network dynamics did not differ between groups (main: t(235)=-1.61, p=0.11; inhibition validation: t(223)=-0.84, p=0.40; shift validation: t(177)=0.57, p=0.57). But interestingly, moment-to-moment SEV and shift network dynamics were more strongly aligned in HCs than in patients in the main dataset (t(235)=2.73, p=0.007; **Figure 4C**). This result was replicated in the inhibition validation dataset (t(223)=-3.25, p=0.001; **Figure 4C**) but not in the shift validation dataset (t(177)=1.59, p=0.11). We further reran analyses after excluding overlapping edges in the SEV and cognitive control networks. The same results were obtained **(Supplementary Materials**), suggesting that temporal alignment existed beyond network spatial overlap, and temporal fluctuations in these brain networks had clinical implications.

## Discussion

Across three independent datasets, we demonstrated a robust association between cognitive control and the ability to recruit recurring brain states flexibly. First, we showed that such flexibility predicted cognitive control in individuals with diverse clinical profiles. Furthermore, within each individual, moments of more flexible brain state engagement occurred alongside moments of network connectivity related to better cognitive control. Altogether, these findings set forth SEV as a new intermediate phenotype underpinning both inter- and intra-individual differences in cognitive control across psychiatric disorders.

### Flexible brain state engagement predicted cognitive control transdiagnostically

Within datasets, patients drove the relationship between SEV and inhibition, while the link between shift and SEV was found in the full sample. It may be possible that questionnaire-based items are more ecologically valid because they often represent real-world scenarios^10,15^. Additionally, HC may be performing near ceiling level for the inhibition task, particularly in the main dataset (**Supplementary Figure 2**). Thus, BRIEF and RRS may have greater power in detecting how the relationship between SEV and cognitive control varies along a continuum in HCs and patients. Higher intrinsic SEV may support cognitive control by enabling more flexible exploration of brain state combinations in response to evolving demands. This interpretation aligned with prior investigations noting the relevance of neural variability for cognition^9,47–49^ and psychopathology^20^. Additionally, our approach aligned with the RDoC framework by moving beyond diagnostic categories^50^. SEV’s ability to capture whole-brain dynamics is additionally advantageous since transdiagnostic features are unlikely to be underpinned by a single brain network^51^.

We next tested whether cognitive control can be inferred from SEV using machine learning. Predictive modeling can strengthen our understanding of psychopathology^52^. Specifically, it helps alleviate concerns about overfitting^53^ and demonstrates the reproducibility and replicability of the observed association^54,55^. Successful external validation illustrated the generalizability of our findings, as the three unharmonized datasets recruited heterogeneous samples and differed in acquisition parameters and measurement tools (color-word vs. Stroop and BRIEF vs. RRS). Replication, despite significant dataset-specific idiosyncrasies, is promising. Patient populations tend to be diverse in real life, varying in symptom manifestations, comorbidities, and treatment histories^56,57^. Models that can successfully predict behaviors across diverse clinical profiles may hold greater potential for explaining brain-behavior relationships in the real world.

We utilized both task-based and questionnaire metrics of cognitive control. Objective and subjective measures can capture different dimensions of cognition^10,15,58,59^. Our findings showed that SEV was highly relevant for multiple aspects of cognitive control. Compared with other behavioral measures, RRS may be more clinically oriented. However, rumination has been linked to cognitive control^60–62^ and spans multiple clinical conditions^63^. We indeed found that models trained and tested with rumination performed well transdiagnostically. Altogether, these results provided preliminary evidence that altered SEV may give rise to aberrant executive functioning, which may, in turn, contribute to emotional dysregulation and greater difficulty disengaging from negative thoughts^23,64^.

### Shared brain networks support flexible brain state engagement transdiagnostically

We further found that shared brain networks predicted SEV across all three transdiagnostic datasets. Predominantly within-network connections gave rise to greater SEV (**Figure 3**). This shared pattern dovetailed with prior results indicating that network integration supported neural variability^65^. Further parallels could be drawn between our findings and literature on network modularity, which described network profiles characterized by more within-than between-module connections^66^. Modularity also supports adaptive behaviors and executive functions, including cognitive control^67–69^. Another particularly relevant observation was that greater within-network connectivity in the DMN and visual networks occurred before hit but not missed task responses^66^. Notably, within-network connectivity in both of these brain networks predicted higher SEV. This overlap suggests that greater network integration may enable behavioral flexibility, with one avenue being supporting flexible brain state engagement.

In all three datasets, lower SEV was associated with stronger connections between the motor and cognitive control networks (**Figure 3**). The motor network has been implicated in substance use as well as cognitive traits relevant for a range of psychiatric conditions^70–72^. Echoing our findings, a data-driven approach has revealed that connections between somatomotor and executive networks underpinned a rich set of behaviors encompassing cognitive processes and clinical symptoms transdiagnostically^73^. One possibility is that stronger connectivity between motor and cognitive control networks may reflect a greater effort in regulating the more automatic behavioral responses supported by the motor network^74^. This network profile may thus represent a brain system that is more prone to giving rise to automatic responses rather than selecting behavioral responses after careful exploration of different brain state combinations.

### Moment-to-moment temporal alignment of SEV and cognitive control networks

Building on prior studies using edge time series to probe behavioral fluctuations over time^46,75,76^, we investigated the temporal alignment of networks predicting SEV and cognitive control. This novel approach is particularly beneficial when evaluating the constructs of interest instantaneously is difficult, as in our case. While we identified brain networks in the main dataset, moments of higher SEV were associated with moments of network connectivity related to better cognitive control across all three datasets. Successful external validation extended our prior findings to demonstrate a generalizable relationship between SEV and cognitive control within the same individual.

We obtained preliminary evidence that compared to HCs, patients showed weaker temporal alignment between SEV and shift network dynamics. Lower alignment could reflect an altered ability in patients to adjust dynamic brain responses to address evolving demands in cognitive control. The largest effect size was observed in the inhibition validation dataset, likely because it focuses specifically on three mental disorders and is more enriched for case-control comparisons. Similarly, we may not be able to detect group differences in the shift validation dataset since it included the lowest number of patients out of the three datasets and a relatively small number of participants diagnosed with each disorder (see clinical diagnosis information in **Supplementary Materials**). Interestingly, we did not observe significant group differences in the temporal alignment between SEV and inhibition. This discrepancy may occur since shift mainly evaluates flexibility in thinking style^15,77^. Temporal alignment was examined during rest, a period characterized by remarkable variations in thought dynamics and thought content^78,79^. Thus, shift, compared to task-based RT measures, may be more behaviorally relevant during this time. As the three datasets did not perform thought sampling during rest, how changes in SEV track thought dynamics should be more formally tested in future work. Additional work with more rapid behavioral samplings is needed to validate whether SEV fluctuates to support behavioral responses to evolving demands.

### Limitations and future directions

This study presented robust evidence demonstrating flexible engagement of recurring brain state was highly relevant for cognitive control in transdiagnostic populations. However, several limitations should be noted. While we aimed to capture cognitive control’s multifaceted nature by including inhibition and shift measures, the extent to which SEV supports different cognitive control subcomponents remains a future research direction. Importantly, different cognitive control tasks may require different operations^4^. Clever task designs are needed to cleanly tease apart how SEV supports the different components of adaptive control. Additionally, while we explored temporal fluctuations in cognitive control and SEV using brain network dynamics, we only examined their relationship during rest. Cognitive control can fluctuate within the same individual under different contexts and demands^1^. Probing whether the relationship between SEV and cognitive control differs in the face of particularly salient stimuli or specific environments^80^ may shed light on whether the transdiagnostic relationship between SEV and cognitive control breaks down in a disorder-specific manner under certain circumstances.

### Summary

Across three independent datasets, we established transdiagnostic associations between flexible brain state engagement and cognitive control. SEV, a multivariate metric approximating how flexible individuals are at recruiting recurring brain states, successfully predicted multiple forms of cognitive control in previously unseen individuals. Additionally, shared brain networks were associated with SEV. Brain network dynamics suggested that moments of greater SEV occurred with moments of network connectivity, reflecting better cognitive control within the same individual. In sum, these findings underscore the promise of SEV as a novel intermediate phenotype for cognitive control across psychiatric conditions.

## Methods

### Datasets

The current study utilized resting-state and task-based fMRI data from three independent, transdiagnostic datasets. Detailed information on the three datasets can be found in prior work^30,37,38^. We briefly summarized characteristics relevant to the current study below. A detailed breakdown of each dataset’s clinical diagnoses is included in the **Supplementary Materials**. Study procedures were approved by the corresponding ethics committee for each dataset.

In the main dataset (i.e., YaleNeuroConnect^30^), clinical diagnoses were determined using the Mini International Neuropsychiatric Interview for DSM-5 and ICD-10 as well as based on self-report mental health or neurological diagnoses. Participants with at least one neuropsychiatric disorder diagnosis were placed in the patient group. Individuals with no known neurological disorder or mental health diagnosis were considered as HCs. From the main dataset, we identified 312 participants with available resting-state and task-based fMRI data. We then excluded 45 participants due to scan or preprocessing issues, 27 for excessive motion (i.e., mean framewise displacement (MFD) over 0.2mm for any scan), and 3 for missing brain coverage. After these exclusion criteria were applied, we retained data from 237 participants, including 92 HCs (age: 31.52±11.99 years old; 51 female/41 male) and 145 patients (age: 31.81±11.24 years old; 81 female/64 male).

In the inhibition validation dataset (i.e., CNP^37^), diagnoses were determined based on the Structured Clinical Interview for DSM-IV and the Adult ADHD Interview. We considered all individuals with a clinical diagnosis as patients. Participants met criteria for HCs if they had no lifetime or current clinical diagnosis. We identified 257 participants with resting-state and task-based fMRI scans. We then excluded 29 participants for having MFD over 0.2mm, 1 for scan or preprocessing issues, and 2 for missing brain coverage. After exclusion, we included 225 participants from this dataset, including 109 HCs (age: 31.07±8.69 years old; 49 female/60 male) and 116 patients (age: 33.90±9.47 years old; 47 female/69 male).

In the shift validation dataset (i.e., TCP^38^), clinical diagnoses were determined by a Structured Clinical Interview for DSM-5. Participants met criteria for HCs if they had no history of clinical diagnosis. Participants with at least one clinical diagnosis were considered patients. Task-based and resting-state data were available from 214 participants. We then excluded 28 participants for excessive motion, 4 for scan or preprocessing issues, 3 for missing brain coverage, and an additional 16 for missing RRS data. We retained 163 participants after exclusion. Out of these individuals, 69 participants met criteria for HC (age: 31.61±13.13 years old; 38 female/31 male) and 94 participants (age: 31.93±12.27 years old; 58 female/34 male/2 others) had at least one clinical diagnosis.

The acquisition parameters for the main, inhibition validation, and shift validation datasets have been detailed elsewhere^30,37,38^. Standard preprocessing procedures^81^ were applied to the structural and functional data from all datasets. In brief, we nonlinearly registered the structural data to the standard MNI-152 space. FMRI data was motion corrected using SPM12 before being linearly aligned to the structural data. We carried out additional data cleaning in BioImage Suite, including regressing covariates of no interest (linear and quadratic drift, a 24-parameter motion model, mean white matter, mean cerebrospinal fluid, and mean gray matter signals). We further temporally smoothed (cutoff frequency approximately at 0.12Hz) the fMRI data before performing parcellation with the Shen-268 atlas^82^ to extract timeseries data.

### SEV extraction and prediction

We evaluated flexibility in recurring brain state engagement using an established framework^27–29^. Specifically, the framework takes in a set of brain states of interest to assess their simultaneous, continuous engagement using non-negative least squares regression (see **Supplementary Figure 1** for an illustration). For each brain state, the regression step returns a coefficient to indicate its engagement at each time point. SEV was operationalized as the standard deviation in continuous engagement over time.

Here, we opted to study brain states extracted from the HCP dataset, as the dataset recruited a large number of participants performing a series of tasks. The rich range of task paradigms enabled the identification of recurring brain activation patterns to support various behaviors spanning from relatively low-level (e.g., motor movement) to relatively high-level (e.g., working memory and emotional processing). Secondly, identifying brain states in the HCP circumvented concerns regarding circular analysis and data leakage.

Following prior literature^39^, we performed dimensionality reduction on these fMRI data using nonlinear manifold learning. We then applied K-means clustering to identify four distinct but recurring brain states. The number of brain states was determined using the Calinski-Harabasz criterion^83^. We then labeled these brain states as fixation, high-cognition, low-cognition, and transition based on what tasks participants were performing during the time points in each cluster. Notably, brain network activation in each brain state aligned with its associated cognitive processes. For instance, the entire motor network was activated during the low-cognition brain state, which included time points where participants were instructed to move different body parts (**Supplementary Table 1**). The fixation brain state, encompassing fixation time points across tasks (**Supplementary Table 1**), showed DMN activation and FPN deactivation (**Supplementary Table 2**). Visual representations of the four brain states are shown in **Supplementary Figure 1**. Given that the HCP dataset recruited healthy individuals, we also reran our analysis using brain states identified from one of the transdiagnostic cohorts. Consistent results were obtained using these new brain states (**Supplementary Materials**). This replication suggests that the flexible engagement of recurring brain states is generally relevant for cognitive control, regardless of the specific brain states examined.

We then sought to predict cognitive control using SEV in previously unseen individuals. SEVs from all four brain states were used as predictors for cognitive control in linear models. A model was first generated in the training dataset using MATLAB *robustfit*. Next, we applied the output coefficients to SEVs from the test dataset to predict each participant’s cognitive control based on their SEV profile. Model performance was evaluated by correlating observed and predicted cognitive control.

### Identification of SEV- and cognitive-control-predictive networks

To investigate what brain networks supported SEV, we leveraged 10-fold CPM^43^ to predict overall resting-state SEV from task-based FC. Given SEV’s close association with cognitive control, we used FC generated from a cognitive control task for prediction (main and inhibition validation datasets: stop signal task; shift validation dataset: Stroop task). Since we were interested in studying the brain networks that underpin SEV and behaviors in a transdiagnostic manner, the full sample (i.e., patients and HCs) was used for training and testing. To limit the number of tests run, we assessed the overall flexibility in brain state engagement by performing PCA on the four SEVs. To avoid data leakage, PCA was performed within each training fold separately, without using any leave-out data during the extraction of the overall PCA. We correlated predicted and observed overall SEVs to evaluate model performance. Only edges that predicted SEV in all 10 folds were included in the final SEV networks. To assess the significance of model performance, we performed permutation testing by rerunning the model after randomly shuffling SEVs across participants. This step was repeated 500 times. P-values were then calculated as the number of times a permutation testing model returned a better prediction than the real model.

We additionally adopted a similar approach to identify cognitive control CPM networks. Specifically, 10-fold CPM was used to predict inhibition and shift in the main dataset. Similar to above, we only included edges that predicted cognitive control in all 10 folds in the final cognitive control networks. Model performance was evaluated by correlating predicted and observed cognitive control. We again determined the significance level using permutation testing.

### Co-fluctuation of SEV and cognitive control network dynamics

We leveraged CPM^43^ and edge time series^44,45^ to investigate whether fluctuations in cognitive control and SEV network dynamics aligned within the same individual. Edge time series is a recently introduced method that can temporally decompose functional connectivity into its moment-to-moment contributions. To compute edge time series for each edge, we first z-scored the two node timeseries before finding the element-wise product. Edge time series was computed for each edge in each CPM network. We then summed across all edges to assess the overall CPM network’s continuous engagement. Since it is difficult to measure inhibition and shift at each fMRI acquisition time point across the three datasets, we leveraged edge time series to compute instantaneous CPM network engagement in order to predict moment-to-moment changes in cognitive control. Temporal alignment was evaluated by correlating SEV and cognitive control network edge timeseries within each participant. Lastly, one-sample t-tests examined whether, across participants, the two networks’ temporal alignment differed significantly from 0. We further performed a two-sample t-test to investigate if their temporal alignment differed between HCs and patients within each transdiagnostic cohort.

