## Supplementary Materials for "Flexible brain state engagement predicts cognitive control transdiagnostically"

Brain state identification

We replicated prior work^1^ to identify recurring brain states. In brief, nonlinear manifold learning and 2-step Diffusion Mapping projected task-based fMRI data from the Human Connectome Project S500 release^2^ into a low-dimensional space. We used data from the minimal HCP preprocessing pipeline. Six tasks (motor, working memory, social, emotional, relational, and gambling) from 390 participants were included for brain state identification^1^. After fMRI data was reduced and projected into a low-dimensional space, time points with similar activation patterns were located closer together. K-means clustering then identified 4 recurring brain states with distinct activation patterns. The number of brain states was determined using the Calinski-Harabasz criterion^3^. After exploring a range of potential state numbers, we found four to have the largest Calinski-Harabasz value (i.e., the ratio between within- and between-cluster dispersion). Based on the dominant task conditions in each brain state (**Supplementary Table 1**), we labeled them as fixation, high-cognition, low-cognition, and transition. The centroid of each state cluster was extracted to be its representative time point for all later analyses.

We performed further characterization by investigating what brain networks were activated and deactivated in each brain state. For each representative time point, we first identified the activated (activation above 0) and deactivated (activation below 0) brain regions in several canonical brain networks. We then computed activation or deactivation percentage by dividing the number of activated or deactivated brain regions by the total number of brain regions in each network (**Supplementary Table 2**).

**Supplementary Table 1.** Number of time points associated with each task condition for each brain state.

|  | Fixation | High-cognition | Low-cognition | Cue/Transition |
| --- | --- | --- | --- | --- |
| Fixation | 635 | 0 | 20 | 65 |
| Cue | 41 | 3 | 6 | 158 |
| Working memory (0 back) | 10 | 56 | 99 | 123 |
| Working memory (2 back) | 1 | 201 | 10 | 76 |
| Emotion (Fear) | 10 | 42 | 0 | 48 |
| Emotion (Neutral) | 23 | 12 | 99 | 16 |
| Gambling (Win) | 0 | 100 | 25 | 35 |
| Gambling (Loss) | 0 | 101 | 10 | 49 |
| Motor (Tongue) | 0 | 1 | 52 | 12 |
| Motor (Left foot) | 8 | 5 | 41 | 13 |
| Motor (Left hand) | 7 | 0 | 45 | 15 |
| Motor (Right foot) | 0 | 0 | 55 | 12 |
| Motor (Right hand) | 0 | 1 | 50 | 15 |
| Social (Mental) | 0 | 113 | 0 | 47 |
| Social (Random) | 0 | 111 | 0 | 109 |
| Relational (Match) | 3 | 9 | 17 | 40 |
| Relational (Relation) | 3 | 90 | 5 | 9 |

**Supplementary Table 2.** Networks showing the highest activation and deactivation percentages for each state.

|  | Fixation | High-cognition | Low-cognition | Cue/transition |
| --- | --- | --- | --- | --- |
| Activation percentages | DMN  (88.89%) | VAs  (100%) | Motor network (100%) | Visual I (100%) |
|  | Motor network (85.71%) | Visual II  (88.87%) | MF  (86.21%) | VAs (72.22%) |
|  | MF  (82.76%) | FP  (82.35%) | Cerebellum  (84%) | Visual II (66.67%) |
| Deactivation percentages | VAs  (94.44%) | Motor network (87.76%) | Visual I  (100%) | MF (89.66%) |
|  | Visual I  (66.67%) | DMN  (83.33%) | Visual II, VAs, and  DMN  (66.67%) | DMN (88.87%) |
|  | Visual II (66.67%) | Subcortical (79.31%) |  | Motor (83.67%) |

Abbreviation: DMN, default mode network; MF, medial frontal network; VAs, visual association network; FP, frontoparietal network.

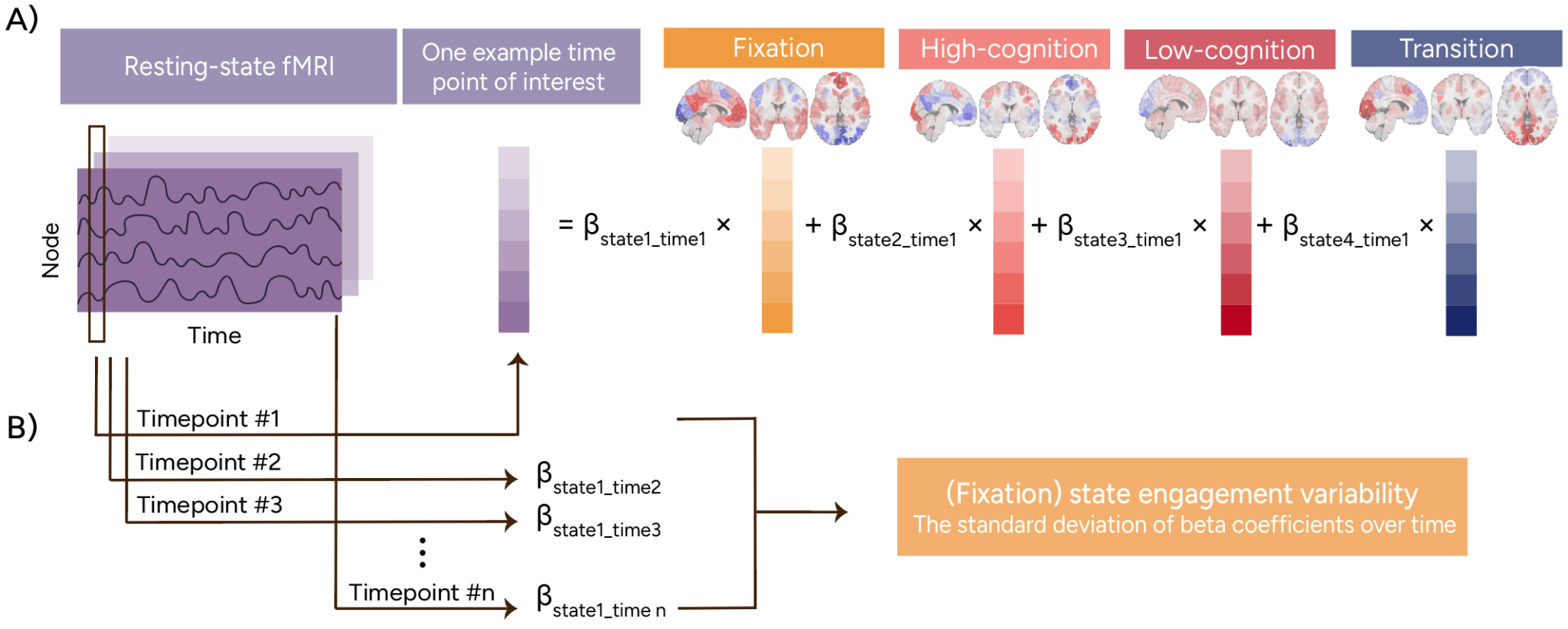

**Supplementary Figure 1. Framework to extract state engagement variability. A)** Our established framework extended recurring brain states identified in the HCP dataset to the three transdiagnostic cohorts. Our framework allows brain state engagement to overlap both temporally and spatially. For each time point, non-negative least squares regression returned a coefficient for each brain state, indicating its contribution at that moment. After extracting moment-to-moment brain state engagement, we computed state engagement variability by computing the standard deviation in continuous engagement over time. **B)** shows the computation of state engagement variability for the fixation brain state as an example. Figure adapted from Ye et al^4^.

**Supplementary Table 3. Relationship between SEV and inhibition within dataset.**

| Overall SEV | Main dataset | Inhibition validation dataset |
| --- | --- | --- |
| All | r=-0.12, p=0.08 | r=-0.27, p<0.01 |
| Patients | r=-0.18, p=0.03 | r=-0.37, p<0.01 |
| HCs | r=0.05, p=0.63 | r=-0.06, p=0.56 |

**Supplementary Table 4. Relationship between SEV and shift within dataset.**

| Overall SEV | Main dataset | Inhibition validation dataset |
| --- | --- | --- |
| All | r=-0.15, p=0.02 | r=-0.16, p=0.04 |
| Patients | r=-0.14, p=0.10 | r=-0.18, p=0.09 |
| HCs | r=-0.07, p=0.51 | r=0.03, p=0.78 |

**Supplementary Table 5. Model parameters used to predict cognitive control in external samples.**

| Model parameters | Fixation | High-cognition | Low-cognition | Transition |
| --- | --- | --- | --- | --- |
| Inhibition model trained in main dataset | -5.47 | -25.67 | 16.85 | 9.73 |
| Inhibition model trained in validation dataset | -528.96 | 113.98 | -111.40 | 245.06 |
| Shift model trained in main dataset | 17.29 | -22.09 | 26.02 | -30.74 |
| Shift model trained in validation dataset | 36.03 | -43.44 | 63.13 | -58.42 |

Replication with brain states identified in a transdiagnostic population

In the main text, we identified brain states in the HCP dataset, which allowed us to identify brain states associated with various behaviors. However, one potential concern that may arise is that since these brain states were identified in mainly healthy individuals, they may introduce bias when being applied to patient populations. While prior literature has shown that similar brain states can be identified in HCs and patients^1,5,6^, we performed additional validation analysis by exploring whether consistent patterns can be observed using brain states generated from one of the transdiagnostic cohorts.

Specifically, we selected the inhibition validation dataset for this purpose. Brain state identification procedures^1^ required time-locked fMRI data where participants were presented with the same stimuli at the same time. The inhibition validation dataset had several fMRI tasks that met this criteria. These tasks included the paired memory encoding, paired memory retrieval, spatial working memory, and task switching paradigms.

Since these paradigms were different from the data we used in the main text, we redid exclusion criteria here again to identify individuals we could use to generate brain states. We first identified 203 participants who completed all four of the fMRI tasks mentioned above before removing 24 participants for excessive motion (mean framewise displacement over 0.2mm for any scan) and 1 participant with missing brain coverage. Out of the 178 participants who met criteria, 102 were patients. To alleviate concerns that the HCP brain states may be more biased towards HCs, we opted to use only data from patients to identify recurring brain states and tested whether we would obtain consistent results. The same nonlinear manifold learning and 2-step diffusion mapping methods were followed to identify brain states. After performing K-means clustering and determining the number of clusters with the Calinski-Harabasz criterion, we identified three recurring brain states with distinctive activation patterns. After identifying the centroid of each cluster, we assessed whole-brain activation similarity between these brain states and the ones identified in HCP using Pearson correlation. We found that brain states identified in a clinical population were highly similar to those from HCP (**Supplementary Table 5**). This pattern aligned with prior literature suggesting that similar brain states can be found in patients and HCs^5,6^.

**Supplementary Table 5. Similarity between inhibition validation and HCP brain states.**

|  | Fixation | High-cognition | Low-cognition | Transition |
| --- | --- | --- | --- | --- |
| Inhibition validation state 1 | r=-0.86; p<0.001 | r=0.65; p<0.001 | r=-0.65; p<0.001 | r=0.87; p<0.001 |
| Inhibition validation state 2 | r=0.86; p<0.001 | r=-0.65; p<0.001 | r=0.63; p<0.001 | r=-0.86; p<0.001 |
| Inhibition validation state 3 | r=-0.68; p<0.001 | r=0.57; p<0.001 | r=-0.13; p=0.04 | r=0.55; p<0.001 |

We then extracted SEV using these new brain states in the main and shift validation datasets. This was not done in the inhibition validation dataset to avoid circular analysis. In the main dataset, overall SEV was again related to inhibition (all: r=-0.17, p<0.01; patients: r=-0.21, p=0.01; HCs: r=-0.07, p=0.50). Similar to what we observed in the main text, the association between SEV and inhibition appeared to be driven by patients here.

However, overall SEV was not linked to shift in the main dataset (all: r=0.03, p=0.69; patients: r=0.05, p=0.57; HCs: r=0.06, p=0.58) or in the shift validation dataset (all: r=-0.04, p=0.57; patients: r=-0.14, p=0.18; HCs: r=0.10, p=0.40). But we were able to use the shift validation dataset to further demonstrate the relationship between cognitive control and SEV. Specifically, we utilized response time during the Stroop incongruent condition in the shift validation dataset. We observed that overall SEV was also correlated with inhibition in the shift validation dataset (all: r=-0.22, p<0.01; patients: r=-0.35, p<0.01; HCs: r=-0.10, p=0.39).

Aligned with our findings in the main text, patients drove the relationship between inhibition and SEV here. Thus, we opted to use data from only patients to further examine whether these newly generated SEVs can predict cognitive control in previously unseen individuals. Since we were only able to establish a relationship between SEV and inhibition within datasets, we did not test the possibility of predicting shift using the new SEVs.

We first trained a model in the main dataset (state 1 beta: 20.43; state 2 beta: -19.31; state 3 beta: -63.97). This model successfully predicted inhibition in the shift validation dataset (r=0.23, p=0.03). Next, we reversed the two datasets and trained a new model in the shift validation dataset to predict inhibition from SEVs (state 1 beta: -76.43; state 2 beta: -303.55; state 3 beta: -136.60). This new model also successfully predicted inhibition in the main dataset (r=0.22, p=0.01).

In summary, these results suggested that SEV is crucial for cognitive control. The replication of our main text results demonstrated that this brain-behavior relationship is unlikely to be driven by biases from using brain states identified in a predominantly healthy population such as HCP. However, these results also indicate that while flexibility in brain state engagement is consistently important for cognitive control, the brain states under investigation matter to some extent. Namely, we were not able to observe an association between shift and the new SEVs. One potential factor contributing to this discrepancy may be the fact that all tasks in the inhibition validation dataset involved high-level cognitive functioning, whereas the HCP dataset encompassed a much richer range of behaviors. Since the shift measures mainly examined flexibility in thinking style, it may be possible that the relative scarcity of fixation time points in the inhibition validation dataset precluded us from capturing brain states sensitive to thinking style.

Spatial overlap in SEV brain networks

To assess whether the three SEV networks shared significant spatial overlap, we followed previous work^7^ and used the hypergeometric cumulative density function to compute the probability of any two networks sharing significant spatial overlap. SEV networks from all three datasets shared significant spatial overlap (**Supplementary Tables 6 & 7**), with the only exception being the positive SEV networks identified from the inhibition validation and shift validation datasets. This may be due to the SEV network identified from the shift validation dataset having a relatively smaller size (**Supplementary Table 8**), potentially contributing to lower spatial overlap with the other SEV networks.

**Supplementary Table 6. Spatial overlap in positive SEV network.**

|  | Main | Inhibition validation | Shift validation |
| --- | --- | --- | --- |
| Main | N/A | p<0.001 | p<0.001 |
| Inhibition validation | p<0.001 | N/A | p=0.29 |
| Shift validation | p<0.001 | p=0.29 | N/A |

**Supplementary Table 7. Spatial overlap in negative SEV network.**

|  | Main | Inhibition validation | Shift validation |
| --- | --- | --- | --- |
| Main | N/A | p=0.005 | p<0.001 |
| Inhibition validation | p=0.005 | N/A | p=0.04 |
| Shift validation | p<0.001 | p=0.04 | N/A |

**Supplementary Table 8. Number of edges in each SEV network**.

|  | Positive SEV network | Negative SEV network |
| --- | --- | --- |
| Main | 263 | 295 |
| Inhibition validation | 165 | 127 |
| Shift validation | 73 | 85 |

Clinical symptom profiles in each dataset

The three transdiagnostic datasets recruited heterogeneous samples. While the inhibition validation dataset focused specifically on three psychiatric conditions, the main and shift validation datasets consisted of transdiagnostic populations with a range of clinical diagnoses. We outlined the clinical breakdown of each dataset below.

In the main dataset, many participants reported comorbidities and may not necessarily have a primary diagnosis. Thus, we reported the number of participants who were diagnosed with each condition here (i.e., some participants may be included in multiple groups below). In the main dataset, 85 individuals were diagnosed with major depressive episode, 63 with generalized anxiety disorder (GAD), 23 with bipolar disorder (BD), 23 with alcohol use disorder (AUD), 26 with substance use disorder, 29 with panic episodes, 25 with past-traumatic stress disorder (PTSD), 12 with social anxiety disorder (SAD), 17 with obsessive-compulsive disorder (OCD), 2 with autism spectrum disorder, 31 with attention-deficit/hyperactive disorder (ADHD), 12 with agoraphobia, 23 with psychotic episodes, and 7 with borderline personality disorder.

In the shift validation dataset, primary diagnosis information was available from each patient participant. We included 3 participants with ADHD, 2 individuals with unspecified anxiety disorder, 1 with binge eating disorder, 7 with bipolar 1 disorder, 4 with bipolar 2 disorder, 1 with cyclothymia, 1 with unspecified depression, 5 with dysthymia, 9 with GAD, 15 with major depression disorder (MDD), 1 with mild cocaine use disorder (CUD), 1 with moderate AUD, 1 with moderate CUD, 1 with OCD, 1 with panic disorder, 1 with past agoraphobia with panic disorder, 3 with past GAD, 11 with past MDD, 1 with past mild AUD, 4 with past PTSD, 1 with past unspecified eating disorder, 1 with premenstrual dysphoric disorder, 7 with PTSD, 1 with severe AUD, 6 with SAD, 4 with schizophrenia, and one patient with missing diagnosis information.

From the inhibition validation dataset, we analyzed data from 37 individuals with schizophrenia, 43 with BD, and 36 with ADHD.

**Consistent results in brain network dynamics were obtained after excluding overlapping edges**

Given SEV and cognitive control’s relevance for behavioral flexibility, it is possible that overlapping edges supported both SEV and cognitive control. Such spatial overlap may bias the examination of these networks’ temporal alignment. Thus, we performed our analyses again after removing overlapping edges. Specifically, we removed 3 overlapping edges from the inhibition positive and SEV negative networks, 9 from inhibition negative and SEV positive networks, 3 from shift positive and SEV negative networks, and 1 from shift negative and SEV positive networks.

All results remained the same after these edges were excluded. Inhibition and SEV network dynamics aligned across all three datasets (main dataset: t(236)=-18.61, p<0.001; inhibition validation dataset: t(224)=-11.05, p<0.001; shift validation dataset: t(178)=-16.34, p<0.001). We also observed temporal alignment in shift and SEV network dynamics (main dataset: t(236)=-12.77, p<0.001; inhibition validation dataset: t(224)=-14.27, p<0.001; shift validation dataset: t(178)=-8.78, p<0.001).

Same as the main text, we found no group differences in the temporal alignment of inhibition and SEV network dynamics (main dataset: t(235)=-1.62, p=0.11; inhibition validation dataset: t(223)=-0.69, p=0.49; shift validation dataset: t(177)=0.48, p=0.63). However, temporal alignment was weaker in patients compared to HCs in the main (t(235)=2.63, p=0.009) and inhibition validation datasets (t(223)=-3.35, p<0.001), but not the shift validation dataset (t(177)=1.58, p=0.11).

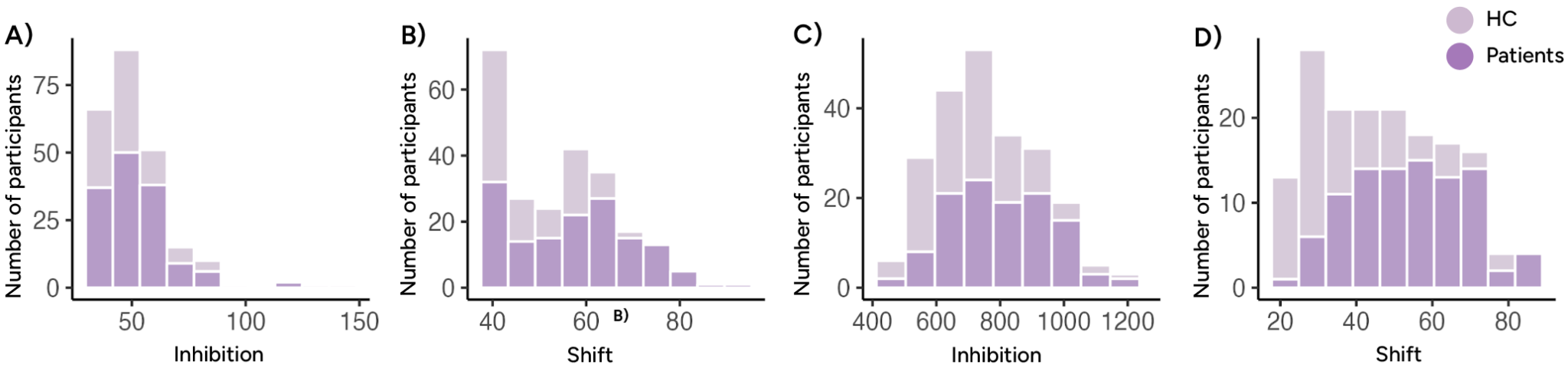

**Supplementary Figure 2.** Cognitive control distributions in the main **A)** and **B)**, inhibition validation **C)**, and shift validation **D)** datasets. Distributions were plotted by group. As shown in **A)**, HC appeared to perform near the ceiling for inhibition in the main dataset. But shift looked more normally distributed in both HC and patients in both the main and shift validation datasets.
